# Regulation of *Sulfolobus acidocaldarius* surface structures by the PP2A core interaction module

**DOI:** 10.64898/2026.08.11.744115

**Authors:** Luis Gayermann, Ankita Banerjee, Shamphavi Sivabalasarma, Friedel Drepper, Pitter Huesgen, Marleen van Wolferen, Sonja-Verena Albers

## Abstract

Protein phosphorylation is a central regulatory mechanism that enables organisms to adapt to changing environmental conditions. The hyperthermophilic archaeon *Sulfolobus acidocaldarius* encodes only two phosphatases: the dual-specificity phosphatase PTP and the serine/threonine phosphatase PP2A. PP2A has previously been implicated in archaellum regulation, and its deletion results in a hypermotile phenotype. Under starvation conditions, PP2A associates with a stress regulatory module comprising the archaellum repressors ArnA and ArnB, the universal stress protein UspA, and a GPN-loop GTPase. Here, we investigated PP2A-associated proteins under normal growth conditions and following UV-induced DNA damage. Pulldown experiments using a genomically HA-tagged PP2A strain identified a PP2A-associated basal regulatory module consisting of ArnA, ArnB, ArnE, and PTP, distinct from the previously described starvation-associated network. In addition, several proteins involved in the biogenesis and regulation of type IV pili co-purified with PP2A. Functional analyses using thermomicroscopy and electron microscopy revealed that deletion of Δ*pp2a*, Δ*arnA*, or Δ*arnB* abolishes Aap-pilus formation and twitching motility, demonstrating that the PP2A regulatory network controls both swimming and surface-associated motility. In contrast, the same network exerted only a modulatory effect on UV-induced cell aggregation. Together, our findings establish PP2A as a central regulator coordinating multiple archaeal surface structures through phosphorylation-dependent signaling.

## Introduction

Phosphorylation is the most abundant post-translational modification (PTM) and serves as a fundamental regulatory mechanism found in all domains of life. It involves the reversible attachment of a phosphoryl group to hydroxyl-containing amino acid residues, thereby regulating protein function, activity, stability and interactions (Hunter 1995, Eichler and Adams 2005, Wang and Wang 2019). In archaea, reversible protein phosphorylation has emerged as an important mechanism for adapting to changing environmental conditions. Archaeal signaling pathways include both two-component and one-component systems; however, members of the Thermoproteota predominantly rely on one-component signaling systems containing serine/threonine/tyrosine kinases (Hanks-type kinases; ePKs) and serine/threonine/tyrosine phosphatases (PPPs). These signaling proteins respond to environmental stimuli and transduce signals within the cell through reversible phosphorylation and dephosphorylation events (Eichler and Adams 2005). Hanks-type kinases are found in all domains of life and are defined by a conserved catalytic core (Esser *et al*. 2016). While ePKs are present across all domains of life, their specific functions in archaea remain poorly understood.

One of the best-characterized outputs of archaeal signal transduction is the regulation of cell surface structures. In the thermoacidophilic archaeon *Sulfolobus acidocaldarius*, interactions with the environment are mediated by different surface structures, each adapted to different environmental conditions. Under nutrient-limited conditions, cells assemble the archaellum, a rotating motility structure that enables swimming toward more favorable environments (Lassak *et al*. 2012, Reimann *et al*. 2012, Nuno De Sousa Machado, Albers, and Daum 2022). In response to DNA damage, cells express UV-inducible pili (Ups pili), which promote species-specific cellular aggregation and facilitate DNA exchange between neighboring cells via the ced-system (Wolferen van *et al*. 2013, 2016, 2020, Recalde *et al*. 2025). In contrast, archaeal adhesive pili (Aap pili) are predominantly expressed during growth in nutrient-rich conditions and contribute to surface attachment, biofilm formation, and twitching motility (Henche *et al*. 2012, Charles-Orszag *et al*. 2024, Gaines *et al*. 2024). In addition, *S. acidocaldarius* produces thread-like surface structures during normal growth, although their physiological function remains unclear (Ng *et al*. 2008, Henche *et al*. 2012, Gaines *et al*. 2022).

The expression of these surface appendages is controlled by a complex regulatory network that integrates environmental signals. Archaellum expression is regulated by several proteins including the regulators ArnR, AbfR1, ArnA, ArnB, as well as protein kinases ArnS, ArnC and ArnD (Reimann *et al*. 2012, Haurat *et al*. 2017, Hoffmann *et al*. 2017, 2019, Li *et al*. 2017, Bischof, Haurat, and Albers 2019). The Lrs14-family protein AbfR1 functions as a phosphorylation-dependent switch between motility and biofilm formation. Phosphorylated AbfR1 promotes extracellular polymeric substance production and biofilm formation while repressing archaellum expression, whereas its unphosphorylated form favors motility (Li *et al*. 2017). The membrane-bound regulator ArnR contains a DNA-binding helix-turn-helix (HTH) domain and regulates *arlB* expression by binding to its promoter. ArnR also binds to promoter regions of genes encoding other type IV pili, including *AapF* and *UpsX*. ArnR1, a paralogue of ArnR, specifically binds the *arlB* promoter under nutrient-limited conditions (Bischof, Haurat, and Albers 2019). DNA damage-induced expression of Ups pili is mediated by the transcription factor Tfb3 and Orc1-2 and is further linked to regulators originally identified in archaellum control, including ArnA, ArnB, and the ArnB paralogue ArnE, highlighting the extensive crosstalk between these signaling pathways (Feng *et al*. 2018, Schult *et al*. 2018, Jiang *et al*. 2023, Liu *et al*. 2025).

The coordination of these regulatory networks is closely linked to phosphorylation-dependent signaling pathways. A phosphoproteomic analysis of *S. acidocaldarius* identified approximately 800 phospho-peptides, indicating extensive phosphorylation-dependent regulation despite the relatively small number of predicted kinases and phosphatases encoded in the genome (Reimann *et al*. 2013). *S. acidocaldarius* encodes eleven predicted typical and atypical Hanks-type kinases but only a single serine/threonine phosphatase, protein phosphatase 2A (PP2A), in addition to one tyrosine phosphatase (Esser *et al*. 2016, Hoffmann *et al*. 2017). This observation suggests that PP2A may function as a major regulatory hub controlling multiple signaling pathways. Consistent with this hypothesis, PP2A has emerged as an important regulator of environmental adaptation among Sulfolobales. Previous studies demonstrated that PP2A influences archaellum expression, cellular motility, and the DNA damage response (Reimann *et al*. 2013, Ye *et al*. 2020, Jiang *et al*. 2023). Furthermore, PP2A was shown to interact with the starvation-induced universal stress protein UspA and with the archaellum regulators ArnA and ArnB, forming a regulatory module that controls motility during nutrient limitation (Ye *et al*. 2020, Ye, Van Der Does, and Albers 2020). These findings suggest that PP2A occupies a central position within phosphorylation-dependent signaling networks governing surface structure expression. Although PP2A has emerged as a key regulator of archaellum expression, it remains unclear whether it also coordinates additional archaeal surface structures through shared phosphorylation-dependent regulatory modules. In this study, we investigated the PP2A interactome during exponential growth and following UV-induced DNA damage to characterize its interaction network under these conditions. Furthermore, we examined the contribution of PP2A-associated proteins to the regulation of the archaellum, archaeal adhesive pili (Aap pili), and UV-inducible pili (Ups pili). Our results reveal a conserved PP2A-centered regulatory module that links phosphorylation-dependent signaling to the coordinated control of multiple surface appendages in *S. acidocaldarius*.

## Results

### PP2A-associated basal regulatory module

To identify proteins associated with PP2A under normal growth conditions, the *S. acidocaldarius* strain MW802, in which the *pp2a* gene is genomically HA-tagged (Ye *et al*. 2020), was used for in vivo co-immunoprecipitation (co-IP) assays with cells harvested at OD600 0.3 (early exponential phase; Fig. 1A) and OD600 0.6–0.8 (late exponential to early stationary phase; Fig. 1B). Enrichment of PP2A-HA was confirmed by Western blot analysis (Fig. S1), and co-purifying proteins were identified by LC-MS. Several previously reported PP2A-associated proteins were enriched with PP2A-HA in both growth phases, including ArnA and ArnB (Fig. 1A, B). Consistent with previous observations during starvation (Ye *et al*. 2020), ArnA and ArnB remained associated with PP2A under both growth conditions, indicating that this interaction extends beyond starvation stress. In contrast, UspA and the GPN-loop GTPase were not significantly enriched under the tested growth conditions, suggesting that their association with PP2A may be restricted to stress conditions (Ye, Van Der Does, and Albers 2020, Korf *et al*. 2023).

**Figure 1.**
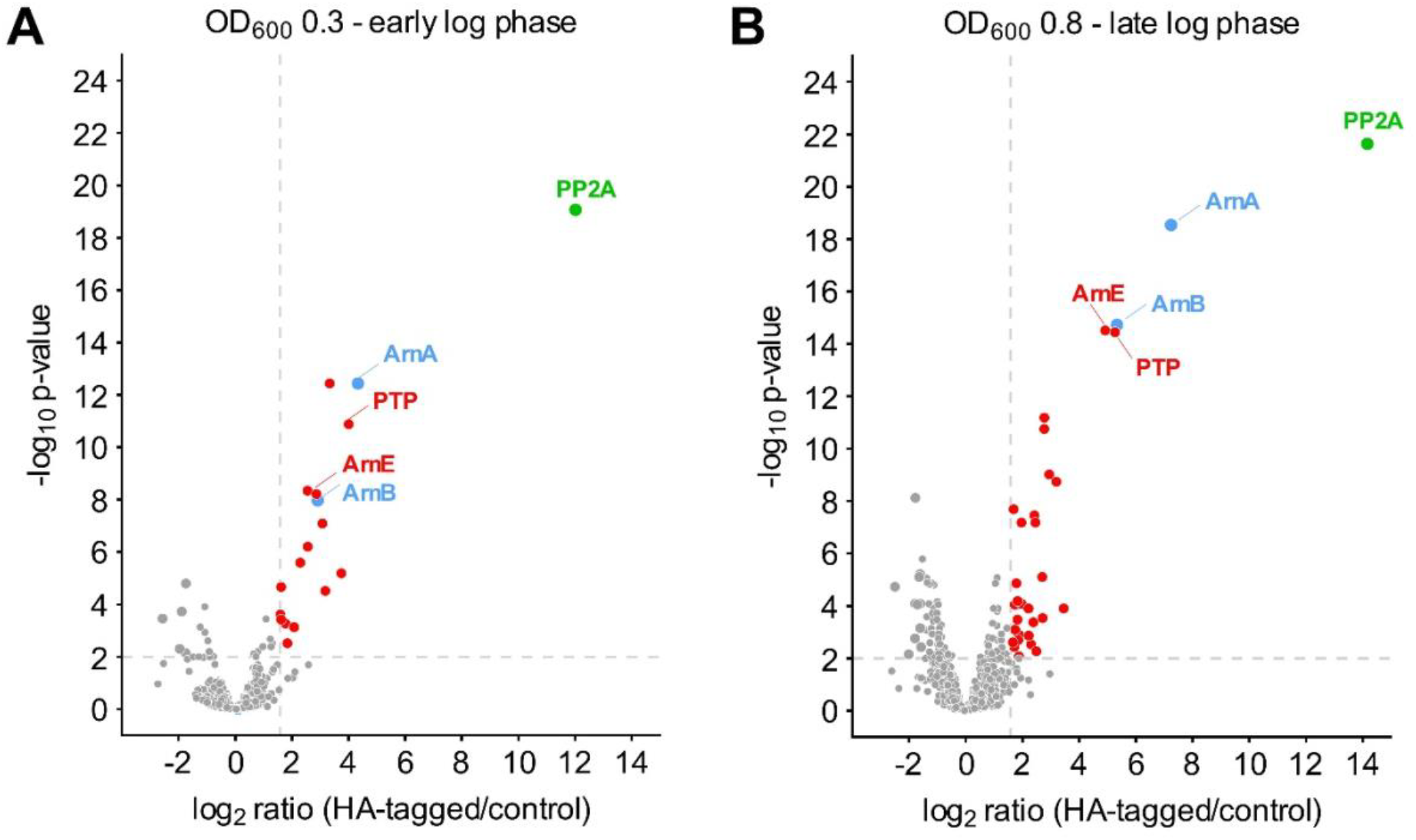
Volcano plots showing interaction partners of the phosphatase PP2A identified by co-IP and LC-MS/MS-based quantitative proteomics. *S. acidocaldarius* strain MW802 expressing HA-tagged PP2A was grown to an OD_600_ of 0.3 (A) or 0.8 (B). Strain MW001 expressing the untagged wild-type PP2A served as the negative control. Each dataset comprises eight biological replicates, processed in two batches of four replicates each. Proteins significantly enriched in the HA-tagged samples (adjusted *p* < 0.01, log_2_ fold change > 1.58) are shown in red, whereas proteins not meeting these thresholds are shown in grey. The bait protein PP2A and its known interaction partners ArnA and ArnB are highlighted in green and blue, respectively. Proteins significantly enriched in the control samples are shown in bigger grey dots. The complete list of identified proteins is provided in Supplementary Table S1.

Interestingly, the dual-specific protein tyrosine phosphatase PTP (Saci_0545) was significantly enriched in both early and late growth phases. As *S. acidocaldarius* encodes only two characterized phosphatases, PP2A and PTP, their consistent co-purification suggests a potential functional relationship. Alternatively, both phosphatases may associate with common phosphorylated substrates. In addition, the ArnB paralogue ArnE (Saci_1209) was enriched under both growth conditions, an interaction not reported before. Previous phosphoproteomic analyses identified phosphorylated serine residues within the N-terminal von Willebrand domain of ArnE, whereas ArnB is phosphorylated at C-terminal threonine residues *in vivo* and *in vitro* (Reimann *et al*. 2013, Hoffmann *et al*. 2017). These distinct phosphorylation patterns suggest that ArnE and ArnB have diverged functionally and may differ in their interactions with ArnA.

Based on the reproducible enrichment of PP2A, ArnA, ArnB, PTP, and ArnE in exponentially growing cells, we define this set of proteins as a “PP2A-associated basal regulatory module”. In contrast to the previously described PP2A stress module, which additionally includes UspA and the GPN-loop GTPase (Ye *et al*. 2020), this module lacks significant enrichment of the latter two proteins under normal growth conditions.

Notably, several proteins encoded by the maltose transporter operon (Saci_1160–1166) were phases. The most strongly enriched protein was the substrate-binding protein MalE (Saci_1165). Although phosphorylation has so far only been reported for the transcriptional regulator MalR (Saci_1161), the consistent enrichment of multiple components of the maltose transport system suggests a previously unrecognized association between PP2A and maltose uptake. A further growth-phase-specific interactor was AapX (Saci_2316), which was strongly enriched during the late growth phase (Fig. 1B). AapX is a component of unknown function of the archaeal adhesion pilus and has previously been identified as a phosphoprotein (Reimann et al., 2013), making it a potential PP2A substrate. Its selective enrichment during the late growth phase suggests that the association of PP2A with AapX is condition-dependent and may contribute to the transition from surface-associated adhesion and twitching motility to archaellum expression and swimming motility.

### The PP2A basal regulatory module mediates twitching motility

To determine whether the PP2A basal regulatory module also affects Aap-pili formation, we analyzed twitching motility in the *S. acidocaldarius* deletion strains Δ*pp2a*, Δ*ptp*, Δ*arnA*, Δ*arnB*, Δ*arnAarnB* and *ΔarnE* by thermomicroscopy at 75°C in the VAHeat system, using the non-twitching strain *ΔaapE* as a negative control. Surface-associated cells were tracked using the ImageJ plugin TrackMate (as described in Charles-Orszag et al. 2024).

As reported previously (Charles-Orszag *et al*. 2024), *ΔaapE* cells frequently detached from the surface and were passively displaced by the flow generated within the VAHEAT chamber. These tracks were characterized by high confinement ratios and strong directional movement. To minimize this bias, tracks representing passive flow-dependent displacement, as well as tracks shorter than 50 frames, were excluded from further analysis (Fig. S2F). Wild-type cells displayed characteristic twitching motility, with cells moving in multiple directions across the surface, as illustrated by the worm plots (Fig. 2A). In contrast, the Δ*pp2a, ΔarnA, ΔarnB*, and *ΔarnAΔarnB* strains resembled the non-twitching *ΔaapE* control strain (Fig. 2B, D–G). These strains exhibited little surface translocation and only occasional short movements, likely due to rapid attachment and detachment events. In contrast, the *Δptp* and *ΔarnE* strains retained twitching behavior comparable to the wild type, displaying non-directional surface-associated movement (Fig. 2A-E, G). Representative detected cells are shown in supplementary movie S1.

**Figure 2.**
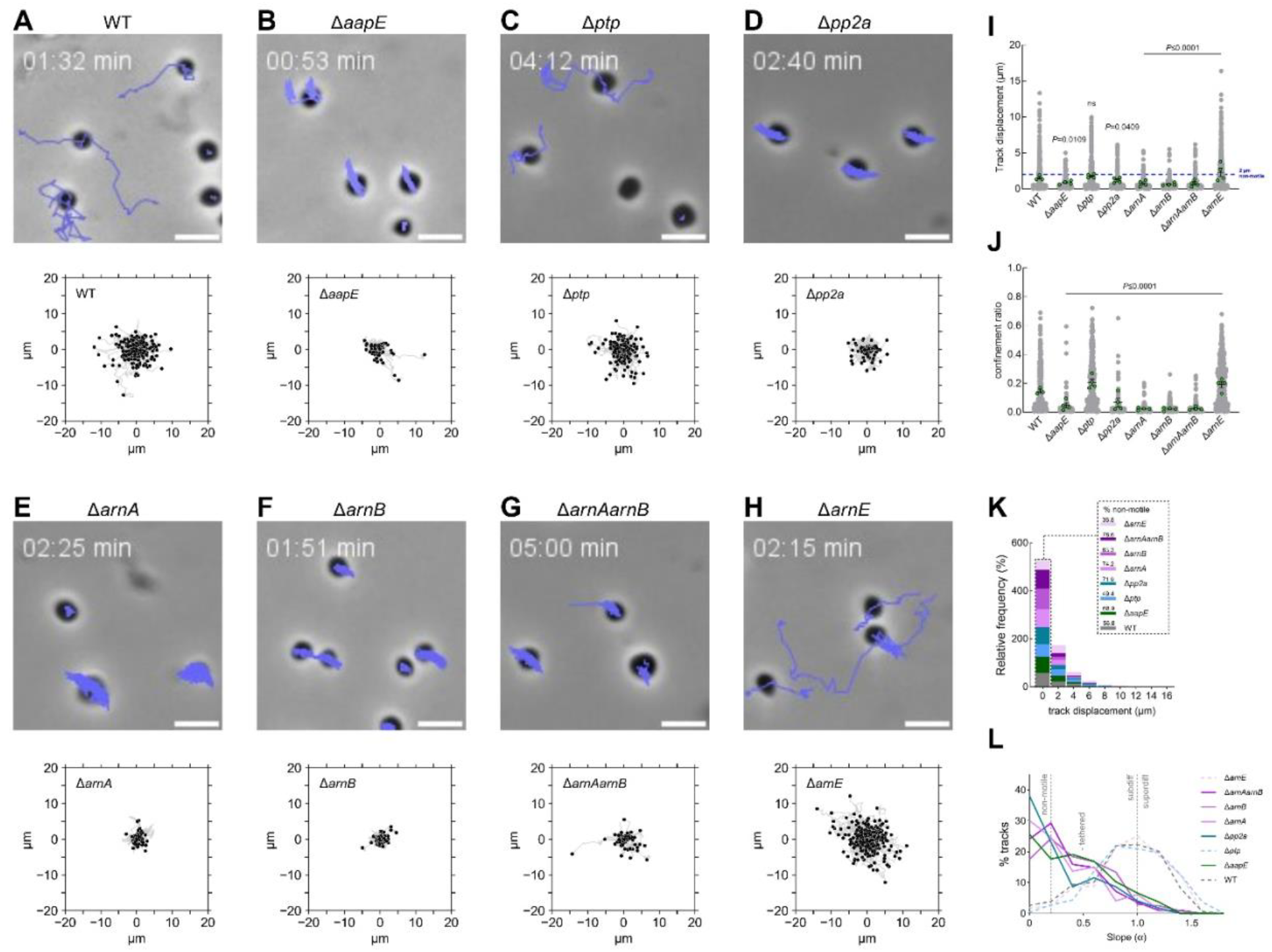
Twitching motility of deletion mutants of the PP2A core regulatory module. Panels (A–H) show representative frames from one of 12 time-lapse movies acquired during live-cell imaging of *S. acidocaldarius at* 75 °C, together with worm plots of all Weka-detected twitching cells for the wild type (492), *ΔaapE* (136), *Δptp* (320), *Δpp2a* (173), *ΔarnA* (146), *ΔarnB* (239), *ΔarnAΔarnB* (195), and *ΔarnE* (509). Quantification of track displacement (I), confinement ratio (J), the relative frequency of twitching cells (K), and the mean squared displacement (MSD) slope (α) (L) is shown for all tracked cells. Tracks shorter than 2 µm were classified as non-motile. In panel (L), dashed lines represent twitching strains, whereas solid lines represent non-twitching strains. Source data are provided in Supplementary Table S2, and a representative twitching movie is available as Supplementary Movie S1.

Quantification of track displacement supported these observations, with cells typically moving >10 µm, whereas tethered cells generally moved <5 µm (Fig. 2I). Wild-type, *Δptp*, and *ΔarnE* cells exhibited substantially greater displacement distances than *Δpp2a, ΔarnA, ΔarnB, ΔarnAΔarnB*, and *ΔaapE* cells. Consistent with their worm plots, *Δptp* cells did not differ significantly from the wild type, whereas all other mutants showed significant changes in displacement. Analysis of confinement ratios further distinguished the strains (Fig. 2J). Although all strains displayed mean values below 0.2, indicating predominantly non-directional movement, *Δptp* and *ΔarnE* cells exhibited significantly higher confinement ratios than the wild type, whereas *Δpp2a, ΔarnA, ΔarnB, ΔarnAΔarnB*, and *ΔaapE* cells showed significantly reduced values. Frequency distributions of track displacement revealed that the majority of cells in all strains were either non-motile or exhibited only limited movement (<2 µm) (Fig. 2K). Nevertheless, wild-type, *Δptp*, and *ΔarnE* populations contained a substantially larger fraction of motile cells than *Δpp2a, ΔarnA, ΔarnB, ΔarnAΔarnB*, and *ΔaapE* strains. Additional motility parameters, including total distance traveled, maximum displacement, and minimum, median, and maximum velocities, supported the same overall grouping of strains (Fig. S2A–E). Notably, the non-twitching strains exhibited greater total distance traveled and higher velocities due to passive movement and continuous tethered cell motion rather than active surface translocation (Fig. S2A).

Mean square displacement (MSD) analysis further confirmed these differences (Fig. S3G). Wild-type, *Δptp*, and *ΔarnE* cells displayed MSD profiles characteristic of confined surface-associated motility, whereas *Δpp2a, ΔarnA, ΔarnB, ΔarnAΔarnB*, and *ΔaapE* exhibited behavior consistent with tethered or non-motile cells. Calculation of log-log MSD slopes yielded values below 1.0 for all strains, indicating sub-diffusive motion. However, Δ*pp2a*, Δ*arnA*, Δ*arnB*, Δ*arnA*Δ*arnB*, and ΔaapE cells consistently displayed slope values below 0.5, a hallmark of tethered or non-motile behavior, whereas wild-type, *Δptp*, and *ΔarnE* cells showed higher values indicative of active twitching motility (Fig. 2L). Together, these results demonstrate that PP2A, ArnA, and ArnB are required for normal twitching motili*ty in S. acidocaldarius*, whereas PTP and ArnE are not required under the conditions tested.

### The PP2A basal regulatory module controls Aap-pilus formation

To determine whether the observed twitching defects were associated with changes in surface appendage morphology, all deletion strains were analyzed by transmission electron microscopy (TEM). Wild-type cells in early exponential phase (OD_600_ 0.4) displayed a dense meshwork of Aap-pili surrounding the cell surface, visible as numerous thin, straight filaments (black arrows, Fig. 3A). As expected, the *ΔaapE* control strain lacked Aap-pili due to the absence of the pilus ATPase (Fig. 3B). Thin filamentous structures, referred to as threads (Henche *et al*. 2012, Gaines *et al*. 2022), were observed in all strains (white arrows, Fig. 3). In agreement with the twitching assays, *Δpp2a, ΔarnA, ΔarnB*, and *ΔarnAΔarnB* cells lacked detectable Aap-vpili (Fig. 3D–G). The absence of these surface appendages is consistent with their inability to perform twitching motility and demonstrates that PP2A, ArnA, and ArnB are required for normal Aap-pilus formation. In contrast, *Δptp* and *ΔarnE* cells displayed surface filaments comparable to those of the wild type (Fig. 3C, H), consistent with their twitching capacity.

**Figure 3.**
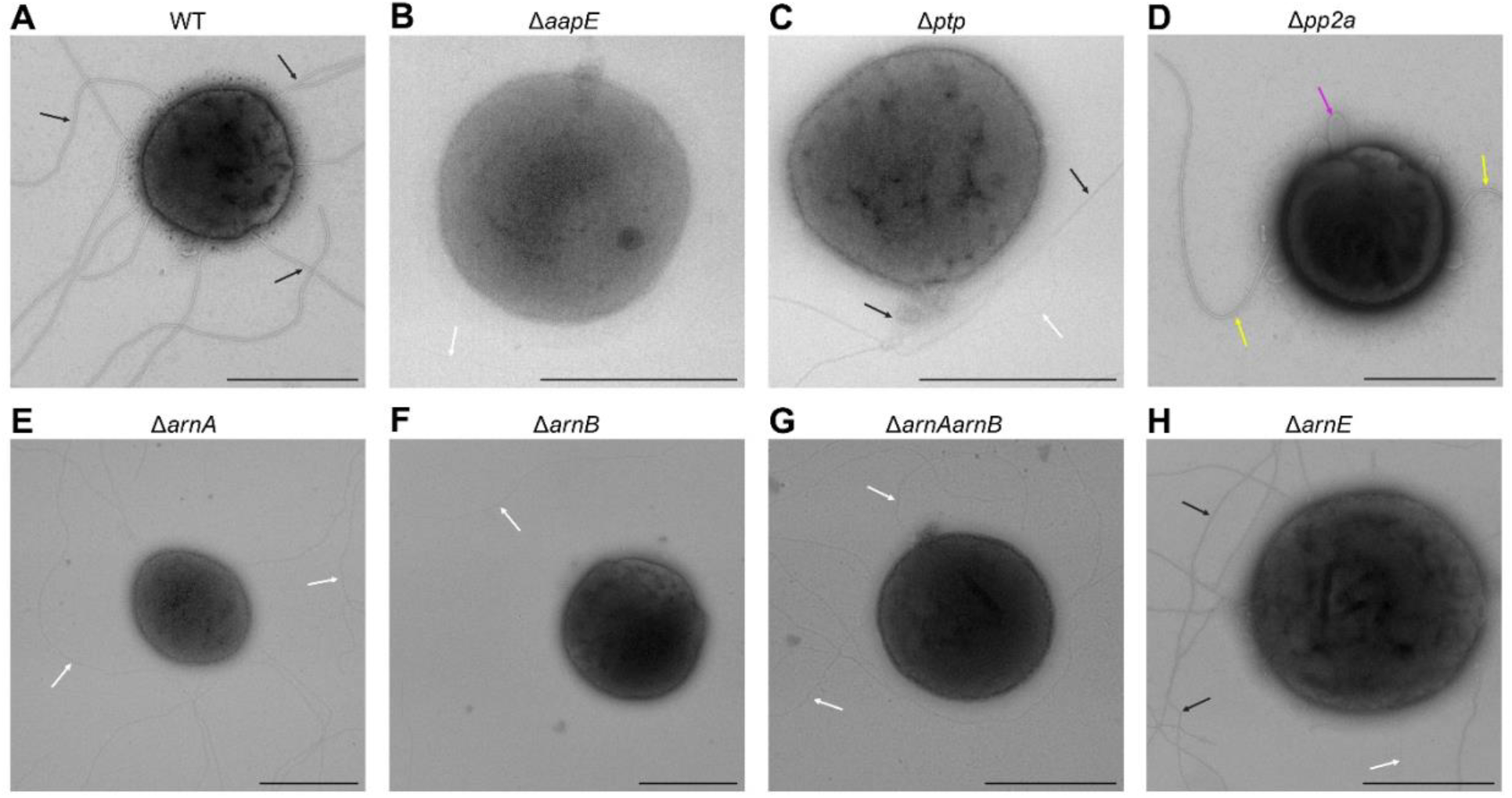
Transmission electron microscopy images of *S. acidocaldarius* wild-type and mutant strains. Panels (A–H) show representative cells of the wild type, *ΔaapE, Δptp, Δpp2a, ΔarnA, ΔarnB, ΔarnAarnB*, and *ΔarnE*, respectively, displaying different surface appendages. Black arrows indicate Aap pili, white arrows indicate thin filamentous structures, yellow arrows indicate archaella, and magenta arrows indicate membrane stress in the *Δpp2a* strain. Scale bars, 1 µm.

Interestingly, a small number of *Δpp2a, ΔarnA*, and*ΔarnB* cells displayed archaella, identifiable by their characteristic curved morphology (yellow arrows, Fig. 3D; Fig. S4A–C). Although observed only rarely, this finding is consistent with the previously described hypermotile phenotypes of these mutants and further supports a role of the PP2A-ArnA-ArnB module in coordinating the expression of distinct surface structures. In addition, a subset of *Δpp2a* cells exhibited pronounced membrane deformations (magenta arrows, Fig. 3D), suggesting that loss of PP2A affects cellular physiology beyond the regulation of motility structures alone. Taken together, the TEM analyses demonstrated that PP2A, ArnA, and ArnB are required for Aap-pilus biogenesis and provide a structural explanation for the twitching defects observed in the corresponding deletion strains.

### The PP2A-associated interactome upon UV irradiation

Besides the archaellum and Aap-pili, *S. acidocaldarius* produces Ups-pili in response to DNA damage. To determine whether UV-induced stress alters the PP2A-associated protein network, PP2A-HA co-immunoprecipitation experiments were performed under non-induced (UV-) and UV-induced (UV+) conditions. Protein enrichment was assessed by comparing PP2A-HA pull-downs with the wild-type control under both conditions, as well as by directly comparing UV- and UV+ PP2A-HA samples (Fig. 4).

**Figure 4.**
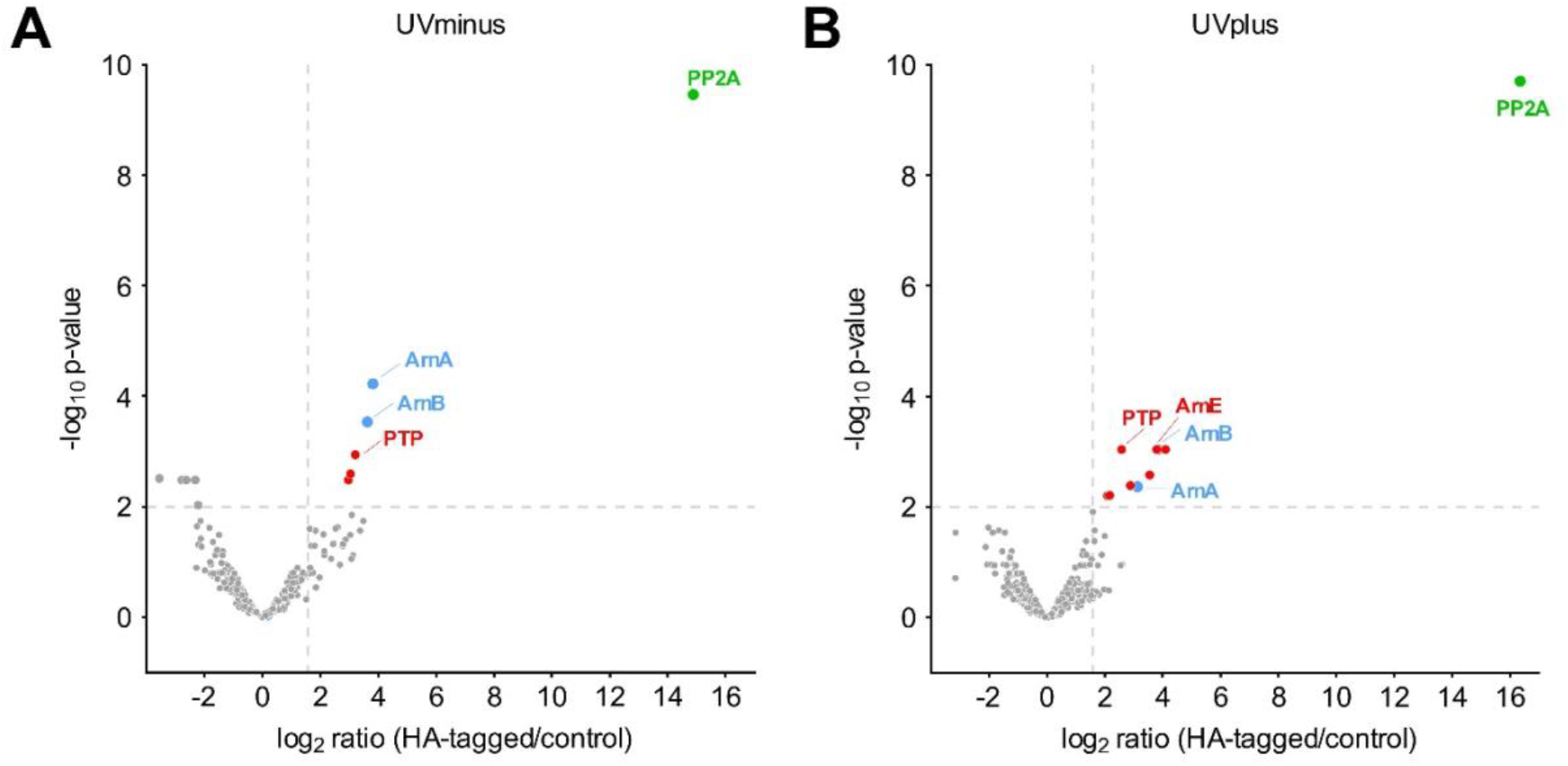
Interaction partners of the phosphatase PP2A identified by Co-IP and LC-MS/MS-based quantitative proteomics following UV irradiation. Volcano plots show proteins enriched in HA-tagged PP2A pulldowns from untreated (UV−) (A) and UV-treated (UV+) (B) *S. acidocaldarius* cultures relative to the untagged wild-type control. Proteins significantly enriched in the HA-tagged samples (adjusted *p* < 0.01, log_2_ fold change > 1.58) are shown in red, whereas proteins not meeting these thresholds are shown in grey. The bait protein PP2A and its known interaction partners ArnA and ArnB are highlighted in green and blue, respectively. Proteins enriched in the control are shown in bigger grey dots. The complete list of identified proteins is provided in Supplementary Table S3.

Under non-induced conditions, PP2A-HA (Saci_0884) was the most strongly enriched protein, confirming efficient immunoprecipitation of the bait (Fig. S1, Fig. 4A). Consistent with the growth-phase experiments, ArnA (Saci_1210), ArnB (Saci_1211), and PTP (Saci_0545) were significantly enriched in the PP2A-HA pull-down. These findings further support the existence of a stable PP2A-associated regulatory module during normal growth.

Upon UV irradiation, the same core set of PP2A-associated proteins was recovered (Fig. 4B). ArnA, ArnB, and PTP remained significantly enriched, and ArnE (Saci_1209) was additionally detected above the significance threshold. UV treatment did therefore not result in the recruitment of a distinct set of PP2A-associated proteins. No proteins were found to be significantly differentially enriched between UV+ and UV-PP2A-HA.

### UV-induced cellular aggregation

To determine whether the PP2A basal regulatory module contributes to the UV-induced aggregation response, aggregation assays were performed with the deletion strains *Δpp2a, Δptp, ΔarnA, ΔarnB, ΔarnAΔarnB*, and *ΔarnE* under non-induced (UV-) and UV-induced (UV+) conditions (Fig. 5, Fig. S4). The *ΔupsE* strain, which is unable to produce Ups-pili and therefore incapable of UV-induced aggregation (Wagner *et al*. 2012, Wolferen van *et al*. 2013), served as a negative control.

**Figure 5.**
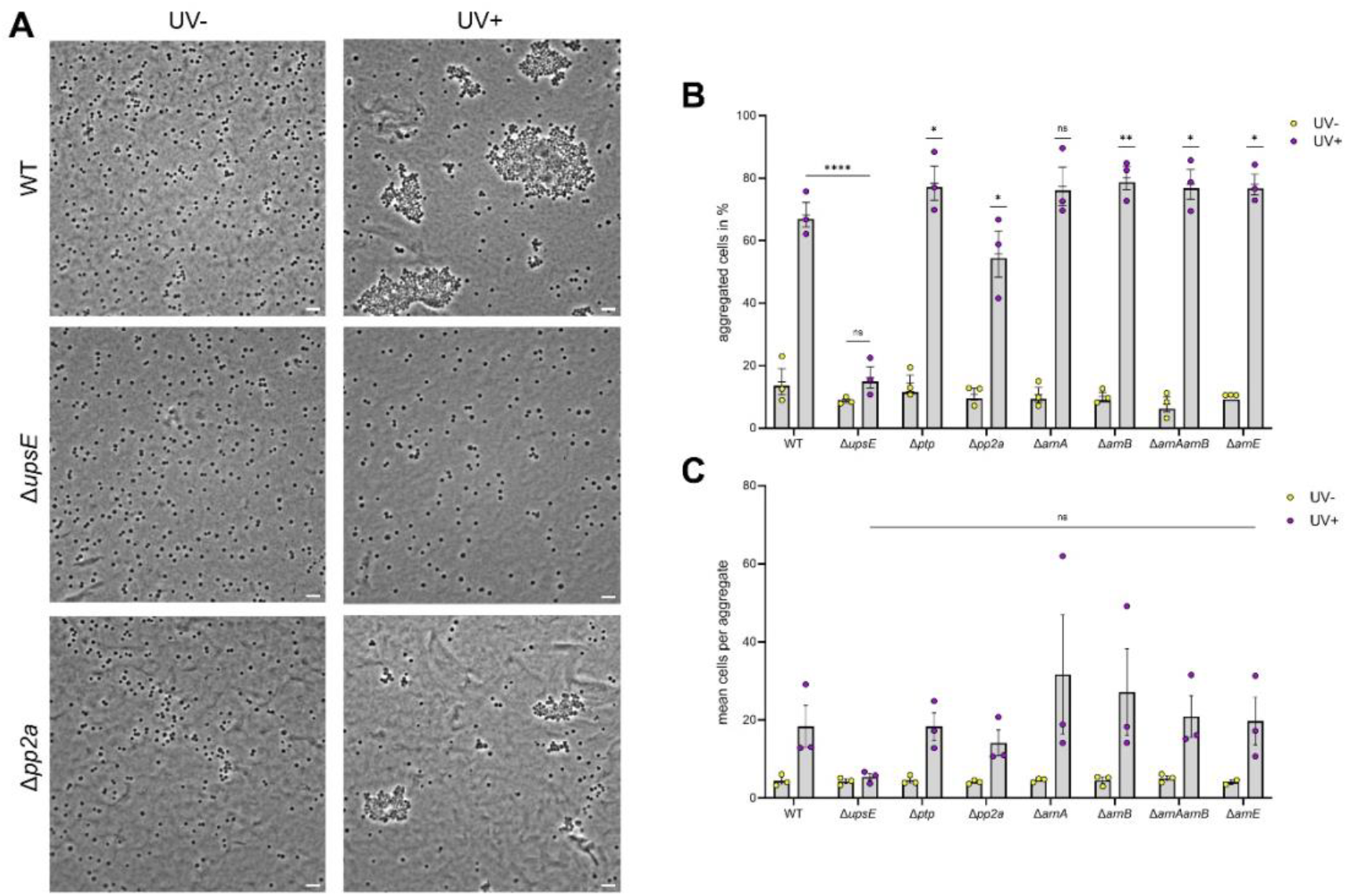
Phase contrast imaging of *S. acidocaldarius* after UV irradiation. (A) Representative images of the wild type (top), the negative control Δ*upsE* (middle), and the Δ*pp2a* mutant (bottom) before UV exposure (UV−; left) and after 3 h recovery following UV irradiation (75 J m^2^; UV+; right). (B) Quantification of UV-induced aggregation based on three biological replicates comprising a total of nine images. Statistical significance was determined using Welch’s t-test by comparing UV-treated wild-type cells with each UV-treated mutant. As expected, the Δ*upsE* control showed no significant difference between UV− and UV+ conditions. (C) Quantification of the average number of cells per aggregate. Statistical significance was determined using Welch’s *t*-test relative to UV-treated wild-type cells.

As expected, wild-type cells exhibited a strong increase in aggregation upon UV irradiation, with the percentage of aggregated cells increasing from approximately 10% under UV-conditions to ~65% under UV+ conditions (Fig. 5B). In contrast, *ΔupsE* showed no significant increase in aggregation following UV treatment, confirming the dependence of this response on Ups-pili. Microscopy images supported these observations, revealing large cellular aggregates in UV-treated wild-type cultures, whereas *ΔupsE* cells remained predominantly dispersed under both conditions (Fig. 5A).

All strains lacking components of the PP2A basal regulatory module retained the ability to aggregate following UV irradiation (Fig. S4). Aggregation frequencies in UV-treated *Δptp, ΔarnA, ΔarnB, ΔarnAΔarnB*, and *ΔarnE* cultures ranged from approximately 75–80%, higher than those observed for the wild type (Fig. 5B). In contrast, *Δpp2a* displayed a modest but significant reduction in aggregation efficiency (~55%) relative to the wild type. Statistical analysis revealed significant differences between all mutant strains and the wild type under UV-induced conditions (Fig. 5B), indicating that deletion of individual module components affects the magnitude of the aggregation response without abolishing it.

Analysis of the mean number of cells per aggregate revealed substantial variability between biological replicates, and no statistically significant differences were detected between strains (Fig. 5C). Nevertheless, *ΔarnA* exhibited particularly high variation in aggregate size, with one replicate containing exceptionally large aggregates comprising more than 60 cells. Although this observation requires further investigation, it suggests that ArnA influences aggregate architecture or stability rather than aggregate formation.

Together, these results demonstrate that PP2A, ArnA, ArnB, ArnE, and PTP are not essential for UV-induced aggregation in *S. acidocaldarius*. However, the altered aggregation phenotypes observed in the mutant strains, particularly the reduced aggregation of *Δpp2a* and the enhanced aggregation of several *arn* mutants, indicate that the PP2A basal regulatory module modulates the aggregation response following DNA damage.

## Discussion

In this study, we identified a PP2A-associated regulatory module present during exponential growth conditions, comprising PP2A, ArnA, ArnB, ArnE, and PTP. Previous work identified ArnA and ArnB as components of a phosphorylation-dependent complex that associates with PP2A during starvation conditions (Ye *et al*. 2020). Our data demonstrate that this association persists during exponential growth, indicating that the PP2A-ArnA-ArnB complex is not restricted to stress conditions, but rather represents a constitutive component of the regulatory module. In addition, we identified ArnE and the dual-specific phosphatase PTP as reproducible PP2A-associated proteins, expanding the previously described PP2A stress module.

In eukaryotes, PP2A activity is controlled through association with a diverse range of regulatory and scaffolding proteins that determine substrate specificity and cellular function (Shi 2009). Although no homologs of the canonical eukaryotic PP2A regulatory subunits have been identified in archaea, the consistent association of PP2A with ArnA, ArnB, ArnE, and PTP suggests that archaeal PP2A likewise operates within stable regulatory protein complexes. Rather than serving as structural homologs of eukaryotic PP2A subunits, these proteins may represent functionally analogous factors that guide PP2A toward specific cellular pathways. The strong conservation of the PP2A catalytic core throughout evolution further supports the idea that diversification of associated regulatory proteins, rather than the phosphatase itself, underlies functional specialization (Kerk *et al*. 2021).

The molecular relationship between PP2A, ArnA, and ArnB has been investigated in detail. PP2A directly dephosphorylates ArnA and ArnB *in vitro* (Hoffmann *et al*. 2017, 2019, Ye *et al*. 2020), suggesting that ArnA and ArnB act primarily as substrates rather than as regulators of PP2A. ArnB interacts with the N-terminal zinc-ribbon domain of ArnA, while phosphorylation of the ArnB C-terminal domain appears to stabilize the ArnA-ArnB complex (Watad *et al*. 2026). Consistent with this model, starvation-induced phosphorylation by kinase ArnC promotes formation of the ArnA-ArnB complex, whereas PP2A-mediated dephosphorylation results in complex dissociation (Ye *et al*. 2020, Watad *et al*. 2026).

This phosphorylation-dependent regulation resembles phospho-switch mechanisms that regulate signaling specificity in eukaryotes. An exemplary mechanism is the DNA damage response during cell cycle checkpoints in *Saccharomyces cerevisiae*, where the FHA-domain of Rad53 recognizes monophosphorylated threonines at Rad53-SCD1, enabling Rad53 activation and downstream signaling during DNA damage (Liao *et al*. 2000, Pike *et al*. 2004, Lee *et al*. 2008). Notably, the FHA domain of the downstream kinase Dun1 preferentially recognizes closely spaced doubly phosphorylated threonines, and increasing the number of phosphorylated residues enhances binding affinity, thereby creating a phospho-counting mechanism that distinguishes different signaling outputs (Lee *et al*. 2008). The phosphorylation state of Rad53 is regulated by PP2C, thus leading to inactivation upon dephosphorylation (Guillemain *et al*. 2007). ArnA is likewise an FHA-domain protein that belongs to the class of di-phosphorylation recognition proteins (Hoffmann *et al*. 2019). Recent work demonstrated that sequential phosphorylation of the ArnB C-terminal domain by kinases ArnD and ArnC generates additional phosphothreonine residues that promote formation of a stable ArnA–ArnB complex (Watad *et al*. 2026). PP2A-mediated dephosphorylation would therefore reset this phospho-switch by destabilizing the complex, analogous to phosphatase-mediated recovery of eukaryotic checkpoint signaling. Furthermore, during starvation, UspA interaction with PP2A enhances phosphatase activity (Ye, Van Der Does, and Albers 2020), which may accelerate this phospho-switch and facilitate the transition between distinct cellular programs, such as the shift between archaellum-dependent swimming and Aap-pilus-dependent surface adhesion/motility.

The identification of ArnE in PP2A pull-down experiments raises intriguing questions regarding the composition and function of Arn-based regulatory complexes. ArnE is a paralogue of ArnB but lacks the conserved C-terminal threonine phosphorylation sites that are essential for ArnB function (Hoffmann *et al*. 2019). The distinct phosphorylation patterns of ArnB and ArnE suggest functional divergence between the two proteins and may influence their interaction with ArnA. In *Sa. islandicus*, it was shown that under UV irradiation the phosphorylation patterns of ArnB and ArnE change (Huang *et al*. 2020). Additionally, ArnA and ArnE are proposed to form an alternative complex by phosphorylation of ArnE and dephosphorylation of ArnB, which is involved in regulating the DNA damage response (Jiang *et al*. 2023). A similar mechanism may exist in *S. acidocaldarius*. However, unlike ArnA and ArnB, deletion of Δ*arnE* did not affect either twitching motility or Aap-pilus formation, indicating that ArnE is not required for these processes under the conditions tested. Thus, the biological function of ArnE remains unresolved and will require further investigation.

The role of PTP within the PP2A regulatory module remains particularly enigmatic. PTP was consistently recovered in PP2A pull-down experiments and represents the only other characterized phosphatase in *S. acidocaldarius*. Because PTP possesses dual-specific phosphatase activity and can target both tyrosine and serine/threonine residues (Tautz, Critton, and Grotegut 2013), it may share substrates with PP2A or function within the same signaling pathway. However, deletion of *ptp* produced no obvious defects in motility, Aap-pilus formation, or UV-induced aggregation, suggesting that PTP does not directly control these processes. Whether the observed association between PP2A and PTP reflects substrate sharing, coordinated phosphor-regulation, or a more direct functional interaction remains unknown.

A major finding of this study is the identification of a previously unrecognized role for the PP2A-ArnA-ArnB module in Aap-pilus-dependent twitching motility. PP2A, ArnA, and ArnB have previously been linked to archaellum regulation and swimming motility, where deletion of any of these genes results in a hypermotile phenotype (Reimann *et al*. 2012, 2013). Here, we demonstrate that these proteins additionally regulate Aap-pilus formation, surface adhesion, and twitching motility. Deletion of Δ*pp2a*, Δ*arnA*, or Δ*arnB* resulted in a complete loss of twitching behavior and the absence of detectable Aap-pili by electron microscopy. These findings identify the PP2A-ArnA-ArnB network as a central regulator of surface-associated behavior in *S. acidocaldarius* (Fig. 6). The observed phenotypes support a model in which the phosphorylation state of the ArnA-ArnB complex contributes to the switch between swimming and surface-associated lifestyles. Such a switch is consistent with recent transcriptomic analyses in *Sa. islandicus*, which revealed cell-cycle-dependent expression of archaella and Aap-pili (Gomez-Raya-Vilanova *et al*. 2025). Here, daughter cells initially express archaella following cell division, whereas Aap-pili predominate during later stages of the cell cycle. The identification of the phosphorylated Aap-pilus-associated protein AapX as a growth-phase-specific PP2A-associated protein is particularly intriguing in this context. Although the molecular function of AapX remains poorly understood, its selective enrichment during late growth phase suggests that PP2A-mediated dephosphorylation may coordinate the transition between swimming and surface-associated motility by regulating components of the Aap machinery.

**Figure 6.**
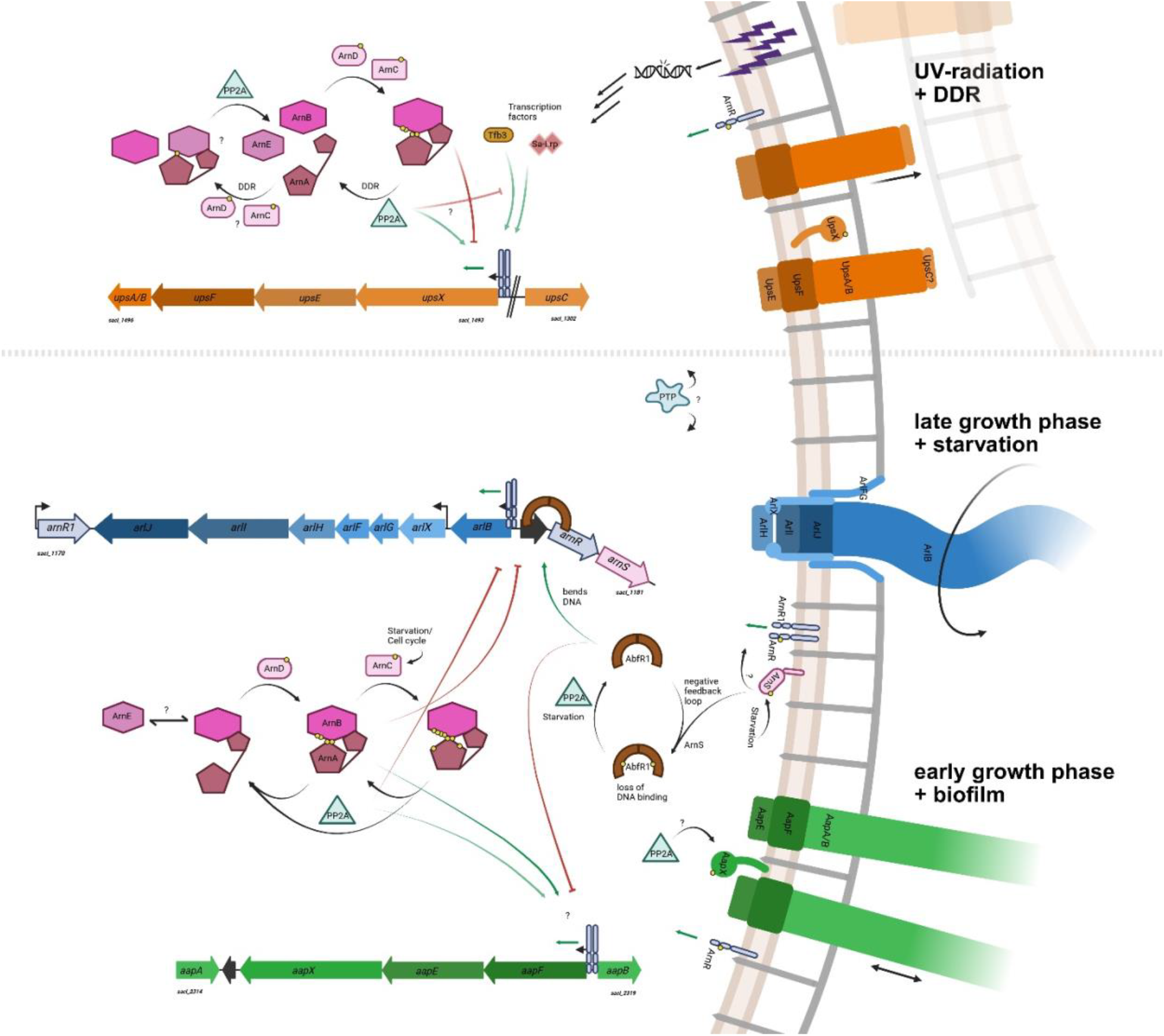
Proposed model for the regulation of surface structures by the PP2A core regulatory module. The model illustrates the three type IV pili-like surface appendages of S. acidocaldarius: the UV-inducible pili (Ups, orange), induced in response to UV irradiation and DNA damage; the archaellum (blue), expressed during starvation and stationary phase; and the archaeal adhesive pili (Aap, green), produced during exponential growth and biofilm formation. The core archaellum regulatory proteins ArnA and ArnB are regulated by reversible phosphorylation mediated by the kinases ArnC and ArnD and the phosphatase PP2A. In response to environmental signals, this core regulatory module coordinates the expression of all three surface appendages. Additional regulators, including AbfR1, Tfb3, and Sa-Lrp, further refine the regulation of individual surface structures.

In contrast to its pronounced role in motility regulation, the PP2A regulatory module appears to play only a minor role during the UV-induced DNA damage response. All mutant strains retained the ability to aggregate following UV irradiation, demonstrating that none of the investigated proteins are essential for Ups-pilus-dependent aggregation. Nevertheless, significant quantitative differences in aggregation efficiency were observed. Deletion of *pp2a* reduced aggregation, whereas deletion of *arnA, arnB, arnAarnB*, or *arnE* resulted in increased aggregation relative to the wild type. These observations are consistent with findings in *Sa. islandicus*, where ArnA, ArnB, and ArnE were proposed to act as secondary regulators that modulate expression of the Ups operon and Ced DNA-transfer system, while key activators such as Orc1-2 and Tfb3 provide the primary regulatory input (Feng *et al*. 2018, Schult *et al*. 2018, Jiang *et al*. 2023, Liu *et al*. 2025).

Interestingly, UV irradiation did not substantially alter the PP2A-associated protein network. ArnA, ArnB, and PTP remained associated with PP2A before and after UV treatment, and no UV-specific interaction partners were identified. These findings suggest that PP2A regulation during the DNA damage response is unlikely to involve major remodeling of the interaction network. Instead, regulation is therefore more likely to occur through dynamic changes in phosphorylation state or protein activity. In this context, phosphorylated DNA damage response regulators such as Tfb3 (Feng *et al*. 2018, Schult *et al*. 2018) or Sa-Lrp (Vassart *et al*. 2013) are attractive candidate PP2A substrates (Fig. 6, upper part).

Taken together, our findings establish a constitutive PP2A-centered regulatory module comprising PP2A, ArnA, ArnB, ArnE, and PTP that regulates multiple cellular processes through a stable protein interaction network rather than through extensive remodeling of its associated factors. Instead, signaling specificity is likely achieved through dynamic changes in substrate phosphorylation and the reversible assembly of phosphorylation-dependent complexes, particularly the ArnA–ArnB module. This regulatory principle resembles eukaryotic PP2A signaling, where stable phosphatase complexes direct signaling through controlled substrate recognition and phosphor-regulation. Functionally, this network acts as a major regulator of motility and surface-associated behavior in *S. acidocaldarius*, controlling both archaellum-dependent swimming and Aap-pilus-dependent twitching motility, while its contribution to the UV-induced DNA damage response appears to be modulatory rather than essential. These findings support a model in which PP2A functions as a central signaling hub that integrates phosphorylation-dependent regulatory pathways governing cellular behavior, environmental adaptation, and stress responses in *S. acidocaldarius*.

## Material and methods

### Strains and Growth conditions of *Sulfolobus acidocaldarius*

*Sulfolobus acidocaldarius* MW001 and derived mutants used in this study (Table S1) were cultivated at 75°C and 120 rpm in Brock basal medium (pH3.0-3.5) supplemented with 0.1% (w/v) NZ-amine, 0.2% (w/v) dextrin and 10 μg/mL uracil (Brock *et al*. 1972, Wagner *et al*. 2012).

**Table 1:** *S. acidocaldarius* strains used.

| Strains | Genotype | Source/Reference |
| --- | --- | --- |
| **MW001** | DSM639 $\Delta pyrE/\Delta pyrF$,<br>uracil auxotroph | Wagner et al.,<br>2012 |
| **MW802** | Chromosomally HA-tagged *saci0884*<br>(*saci_pp2a*) gene at the<br>C-terminus | Ye et al., 2020 |
| **MW025** | MW001 $\Delta pp2a$<br>($\Delta saci\_0884$) | Reimann et al.,<br>2013 |
| **MW351** | MW001 $\Delta arnA$<br>($\Delta saci\_1210$) | Reimann et al.,<br>2012 |
| **MW353** | MW001 $\Delta arnB$<br>($\Delta saci\_1211$) | Reimann et al.,<br>2012 |
| **MW356** | MW001 $\Delta arnE$ ($\Delta vWA2$)<br>($\Delta saci\_1209$) | Reimann et al.,<br>2012 |
| **MW376** | MW001 $\Delta arnAarnB$<br>($\Delta saci\_1210saci1211$) | Hoffmann et al.,<br>2019 |
| **MW010** | MW001 $\Delta ptp$<br>($\Delta saci\_0545$) | Reimann et al.,<br>2013 |
| **MW160** | MW001 $\Delta aapE$<br>($\Delta saci\_2318$) | Henche et al., 2012 |
| **MW109** | MW001 $\Delta upsE$<br>($\Delta saci\_1494$) | Wagner et al.,<br>2012 |

### HA-Pulldown using magnetic beads

*Sulfolobus acidocaldarius* strains MW802 (endogenously HA-tagged PP2A) and MW001 (background strain) were each grown in 400 mL medium and subjected to different treatments. For growth phase comparisons, cultures were harvested at an OD_600_ of 0.3 (early growth phase) or 0.8 (late growth phase). For UV-induced pulldown experiments, cultures were grown to an OD_600_ of 0.3 and exposed to UV-light (Spectrolinker™ XL-1000 UV Crosslinker, Spectronics Corporation) with 75 J m^-2^ (Fröls *et al*. 2007, Wolferen van *et al*. 2013). The 400mL cultures were split into 200mL UV-induced (UV+) and 200mL UV-non-induced (UV-) as a control. Both conditions were recovered at 75°C, 120 rpm for 30 min. The pulldown procedure was kept the same as the normal growth pulldown

For each co-IP assay (adapted from Ye *et al*. 2020), 400 mL of *S. acidocaldarius* culture was used. The culture was cooled down on ice until the temperature approached 37°C, and cells were collected by centrifugation at 4648*g (Anvanti J-26 XP Beckman Coulter, JLA10.500 rotor), 4°C for 20 min. For cell lysis, the pellet was resuspended in lysis buffer (25 mM Tris, pH 7, 150 mM KCl, 10 mM EDTA, 5% glycerol) supplemented with protease inhibitor (Thermo Fisher) and DNase I (Roche) to an OD_600_ of 40. Cells were disrupted by solubilization using a final concentration of 2% DDM at 37°C while rotating for 1h. Cell debris was removed by centrifugation at 4863*g at 4°C for 20 min (ROTINA 380R, Hettich) and subsequently at 200,000*g and 4°C for 1h (Optima™ MAX-XP Ultracentrifuge, Beckman Coulter). After centrifugation, the supernatant was used for co-IP assays with Pierce HA-tag IP/co-IP kit (Pierce) as described in the manufacturer’s instructions. Lysate, Flowthrough, three wash steps and 25 μL of the elution fraction was used for SDS-PAGE. The proteins in the co-IP elution fractions were separated and blotted to PVDF membrane (Roche). The membrane was incubated with primary antibody against the rabbit anti-HA-tag (Sigma). Subsequently, secondary goat anti-rabbit-HRP antibody (Invitrogen) was used. Chemiluminescent signals were visualized by the ECL Chemocam Imager (INTAS) with the Clarity Western ECL blotting substrate (Bio-Rad). Additionally, the co-IP elution fraction was analyzed by mass spectrometry.

### LC-MS analysis and data processing

The elution fractions from the co-IP experiments were denatured in 1% SDS and cysteine residues were reduced with 10 mM DTT for 15 min at 95°C. Free thiol groups were alkylated using 50 mM chloroacetamide for 30 min at 22°C. The reaction was quenched with 50 mM DTT. For tryptic digestion, SP3 bead-based purification protocol (Hughes *et al*. 2019) was used. Proteins were absorbed to the beads (Sera-Mag SpeedBead magnetic carboxylate-modified particles, Cytiva) with a 1:10 final protein:bead (w:w) ratio in 80% ethanol for 5 min at 24°C and 1000 rpm. The samples were three times washed with 80% ethanol. Digestion was performed in 50 µl of 100 mM ammonium bicarbonate containing 0.3 µg trypsin (Promega V5111, Madison, USA) over night at 37°C and 1000 rpm and was stopped by adding 18 µl of 2% TFA. Samples from UV irradiation experiments were prepared similarly, except that the purification, and the tryptic digestion were performed on a Hamilton STARlet automated liquid handling system (Hamilton) using a custom SP3 purification protocol (Joest *et al*. 2026). Peptide mixtures were desalted using self-packed SDB-RPS Stop And Go Extraction tips as described (Joest *et al*. 2026) and finally reconstituted in 0.1% TFA for direct analysis by LC-MS/MS.

For LC-MS analysis, an UltiMate™ 3000 RSLCnano system was online coupled to an Orbitrap Exploris 480 mass spectrometer (both Thermo Fisher Scientific, Dreieich, Germany). The peptide mixture was washed and preconcentrated on a μPAC™ C18 trapping column (PharmaFluidics) with a flow rate of 10 μL/min and peptides were separated on a μPAC™ C18 pillar array column (50 cm bed length, PharmaFluidics) using a binary buffer system consisting of A (0.1% formic acid) and B (86% acetonitrile, 0.1% formic acid). With a flow rate of 0.5 μL/min peptides were eluted applying a linear gradient from 1 to 10% B in 21 min, 10 to 24% in 42 min, 24 to 31% in 13 min, 31 to 47% in 11 min and 95% B for 8 min. For electrospray ionization of peptides, a Nanospray Flex ion source (Thermo Fisher Scientific) with a μPAC Flex iON Connect ESI-MS interface (Thermo Fisher Scientific, Dreieich, Germany) and a fused silica emitter (20 μm inner diameter, 36 μm outer diameter, MicrOmics Technologies LLC) with a source voltage of +1,800 V and an ion transfer tube temperature of 280 °C was used. Cycles of data-independent acquisition (DIA) consisted of: one overview spectrum (RF lens of 40%, normalized AGC target of 300%, maximum injection time of 45 ms, m/z range of 350 to 1,400, resolution of 120,000) recorded in profile mode followed by MS2 fragment spectra (RF lens of 50%, normalized AGC target of 1000%, maximum injection time of 54 ms) generated sequentially by higher-energy collision-induced dissociation (HCD) at a normalized energy of 28% in a precursor m/z range from 361 to 450 in 6 windows of 14 m/z isolation width, in a precursor m/z range from 450 to 800 in 50 windows of m/z isolation width and in a precursor m/z range from 800 to 1,100 in 21 windows of 14 m/z isolation width, overlapping by 1 m/z and recorded at a resolution of 30,000 in profile mode. Samples from UV irradiation experiments were analysed similarly, except that a Q Exactive mass spectrometer (Thermo Fisher Scientific, Dreieich, Germany) was used in DIA mode as described (Joest *et al*. 2026).

MS raw data were converted to mzML format using the ProteoWizard software, version 3.0.21229 (Chambers *et al*. 2012). For protein and peptide identification and quantification a library-free search was performed using DIA-NN version 1.8.1 (Demichev *et al*. 2020) against the UniProt proteome set for *Sulfolobus acidocaldarius* (taxonomy ID 330779, database version 2023-02-27). Trypsin was set as protease with 2 allowed missed cleavages. Carbamidomethylation of cysteine was selected as fixed modification, N-terminal excision of methionine and oxidation of methionine were selected as variable modifications with a maximum number of 2. For precursor ion generation, peptides ranging from 7 to 35 amino acids, precursor charge from 2 to 4 precursor *m/z* range from 350 to 1,100 and fragment ion *m/z* range from 200 to 2,000 were used. Further parameters were: match between runs, unrelated runs, deactivated heuristic protein inference, robust LC quantification. For the analysis of samples from UV irradiation experiments, the search was performed using FragPipe, version 22.0 with quantification by Dia-NN, version 1.82 as described (Joest *et al*. 2026). The protein group quantities output files from DIA-NN were used and all proteins with non-zero label-free quantification (LFQ) intensities in 4 out of 4 replicates in either control or pulldown group were selected for further analysis. Missing values were imputed in two steps: proteins with 3 missing values in one group (missing completely) were subjected to left-censored imputation using the MinProb method, followed by sequential imputation using the R package impSeqRob for proteins with ≥1 remaining missing values. For differential abundance analysis, log_10_ transformed LFQ intensities were subjected to a moderated t-test using linear modelling implemented in the R package limma (Ritchie *et al*. 2015) with correction of p-values for multiple testing. To account for technical variation from processing series of biological replicates on different days, an additional categorical factor for batch correction was included in the linear model design matrix.

### UV Aggregation assay

For the aggregation assay, the deletion strains of *ΔarnA, ΔarnB, ΔarnAarnB, ΔarnE, Δpp2a, Δptp and ΔupsE* were grown to an OD_600_ of 0.2 in a 50 mL culture and exposed to UV-light (Spectrolinker™ XL-1000 UV Crosslinker, Spectronics Corporation) with 75 J m^-2^ (Fröls *et al*. 2007, Wolferen van *et al*. 2013). Half (25 mL) of the culture was transferred to a Petri dish and irradiated with UV light. After treatment, the cultures were recovered at 75°C, 120 rpm for 3 hours. The remaining 25mL culture served as a non-induced control. To determine the aggregation of the different deletion mutants, the cells were diluted to a theoretical OD of 0.2 for comparison. 5µL of the dilution was plated on pads of 2% agarose with Brock basic medium supplemented with NZ-amine, dextrin, and uracil. After drying the cells on the agarose pads, a coverslip was placed. The images were captured using phase-contrast microscopy. Three images were taken per strain and repeated in biological triplicate. Free and aggregated cells (>3) were counted against the total number of cells, as well as the number of aggregates per image. The percentages of cells in aggregates and the average aggregate size were subsequently calculated.

### Negative stain transmission electron microscopy

A 5 µL suspension of *S. acidocaldarius* cells was applied to freshly glow-discharged 300 mesh carbon-coated copper grids (Plano GmbH, Wetzlar, Germany), followed by an incubation period of 30 s at RT. Excess liquid was blotted away, and cells were re-applied on the grid. This was repeated three times. Grids were stained with 2% Uranyl acetate. A Hitachi HT8600 transmission electron microscope (TEM) operated at 100 kV and equipped with an EMSIS XAROSA CMOS camera was used for imaging.

### Twitching assay

Cultures were only grown until an OD_600_ of 0.5 to reduce the number of cells in order to simplify the tracking of cells. For live cell imaging, the VAHEAT (Interherance, Erlangen) was used to image cells at 75°C. In a small (~880 µL) heated chamber the different strains were captured: three times, for 5 min with 1 frame per second (300 frames). The experiment was carried out in biological quadruplicates. Movies were evaluated by following similarly the procedure in Charles-Orszag et al. 2024. The first four frames from a representative movie of MW001 were used to train the Weka detector (Arganda-Carreras *et al*. 2017) in Fiji (Schindelin *et al*. 2012) after decreasing the frame size to 1104×1104 pixels. Weka was trained to detect surface-attached cells (in focus) and ignore unattached cells (out of focus) as well as signals from the background. These categories were uploaded in TrackMate7 (Ershov *et al*. 2022) operated in Fiji and cells were automatically tracked in full-length movies. Low-quality tracks, tracks corresponding to multiple cells wrongfully detected as one, or tracks including cell-cell interaction events were discarded by adjusting the detection threshold in TrackMate7 and manual pruning. The TrackMate7 included LAP Tracker (Frame to Frame 5 microns; Max. Track Gap Closing (5 microns distance, 3 microns gap) was used to track cells. VAHEATs electric current creates a flow in which cells are flowing along, often resulting in long tracks if the cells are not properly attached to the surface. These long flow tracks were manually filtered out by adjusting linearity of forward progression and number of spots in track.

## Data availability

The mass spectrometry proteomics data have been deposited to the ProteomeXchange Consortium via the PRIDE (Perez-Riverol *et al*. 2025) partner repository with the dataset identifier PXD081991.

## Authors contributions

**LG:** Conceptualization, Data curation, Formal analysis, Investigation, Methodology, Validation, Visualization, Writing – original draft, Writing – review & editing; **AB:** Investigation, Methodology, Visualization; **ShS:** Investigation, Methodology, Visualization; **FD:** Data curation, Methodology, Visualization; **PH:** Funding acquisition, Methodology; **MvW:** Conceptualization, Formal analysis, Investigation, Methodology, Supervision, Validation, Writing – review & editing; **SVA:** Conceptualization, Funding acquisition, Methodology, Supervision, Project administration, Resources, Validation, Writing – review & editing

## Acknowledgements

We thank Bettina Knapp for sample preparation for MS analysis. LG was funded by the European Research Council (ERC; Grant: ARCHCELLORG, 2100219401). Views and opinions expressed are, however, those of the author(s) only and do not necessarily reflect those of the European Union or the European Research Council. Neither the European Union nor the granting authority can be held responsible for them. AB was funded by the Marie Skłodowska-Curie Actions through the ARCTECH consortium (Grant No. 21005010901). ShS, FD and PH received funding from the German Research Foundation (DFG) through Collaborative Research Centre SFB 1381 (Project No. 403222702 – SFB 1381). MvW received funding from the DFG under Germany’s Excellence Strategy (CIBSS – EXC-2189, Project ID 390939984). In addition, S-VA was supported by the Volkswagen Foundation (Momentum Grant: Az-94993) and the European Research Council (ERC; Grant: ARCHCELLORG, 2100219401).

