## Supplementary figures for "Regulation of *Sulfolobus acidocaldarius* surface structures by the PP2A core interaction module"

*First author

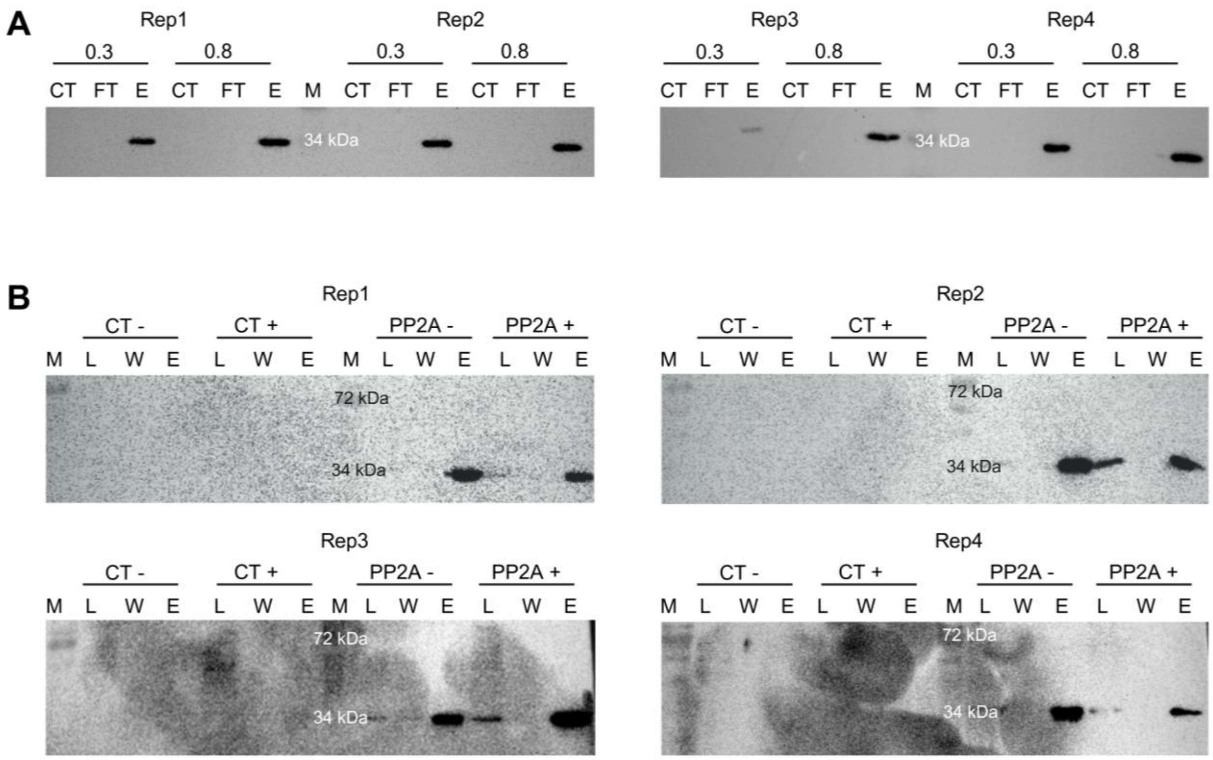


**Figure S1 Western blot analysis of PP2A-HA Co-IP experiments in *S. acidocaldarius.*** Co-IP was performed using anti-HA magnetic beads with the HA-tagged PP2A strain, while the wild-type served as a negative control. SDS-PAGE samples included the cell lysate (L), flowthrough (FT), wash fraction (W), and eluate (E). Western blots were loaded with the wild-type eluate (WT), PP2A-HA flowthrough (FT), and the final PP2A-HA eluate (E). (A) Co-IP experiments performed with cultures harvested at $OD_{600}$ 0.3 and 0.6–0.8. One representative set of four biological replicates is shown. (B) Co-IP experiments performed with untreated (UV−) and UV-treated (UV+) cultures. Membranes were probed with primary anti-HA (rabbit) and secondary HRP-conjugated anti-rabbit (goat) antibodies; therefore, only HA-tagged PP2A is detected, with a signal expected only in the eluate.

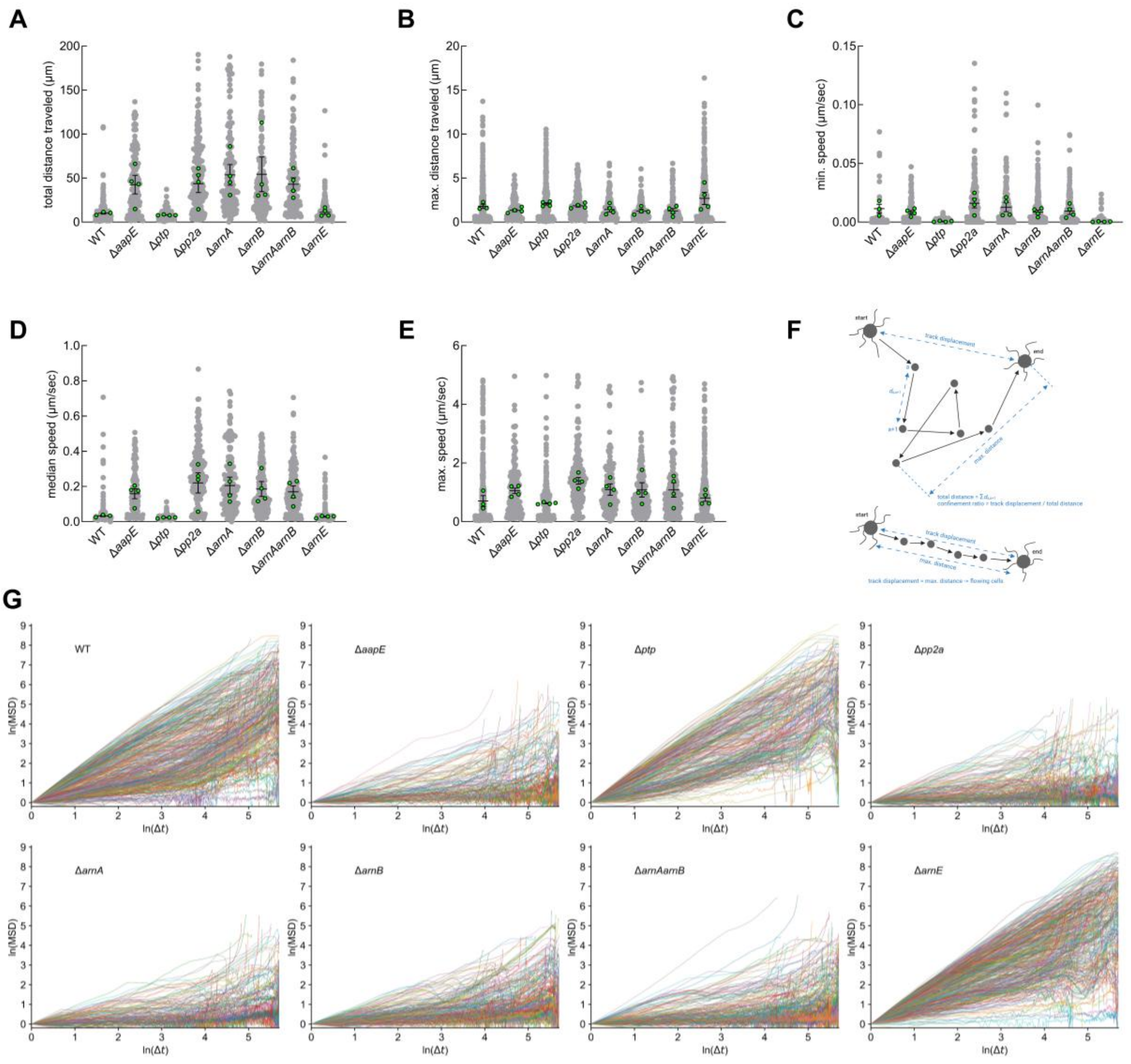


**Figure S2. Additional analysis of twitching motility in *S. acidocaldarius* wild-type and deletion strains.** Panels (A–E) show the total distance travelled (A), maximum distance travelled (B), minimum speed (C), median speed (D), and maximum speed (E) of all tracked cells for each strain. Green dots represent individual biological replicates (n = 4), each comprising three analysed videos. Panel (F) illustrates the filtering strategy used to exclude flowing or floating cells based on identical values for track displacement and maximum distance travelled. Panel (G) shows the mean squared displacement (MSD) analysis of each strain. Twitching strains (wild type, *Δptp*, and *ΔarnE*) display greater mean displacement over time than non-twitching strains (*ΔaapE*, *Δpp2a*, *ΔarnA*, *ΔarnB*, and *ΔarnA* arnB). Negative displacement values indicate movement back towards the starting position. Source data are provided in Supplementary Table S2.

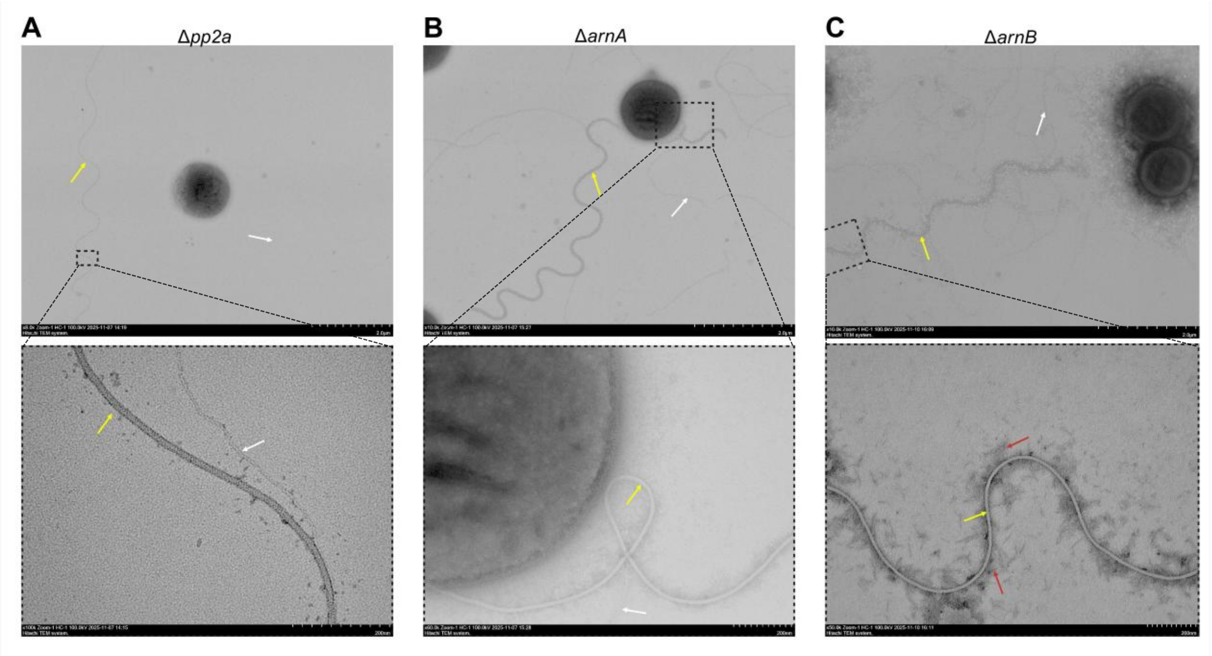


**Figure S3 Archaella in *S. acidocaldarius* deletion strains.** Deletion strains lacking twitching motility are hypermotile on semi-solid motility plates, which mimic starvation conditions that induce archaellum expression. Transmission electron microscopy of exponentially growing cultures ($OD_{600}$ 0.4), used for the twitching motility assays, revealed occasional archaella in these strains (A–C). The lower panels show magnified views of the boxed regions in the corresponding upper panels. Yellow arrows indicate archaella, white arrows indicate filamentous structures, and red arrows indicate staining artefacts.

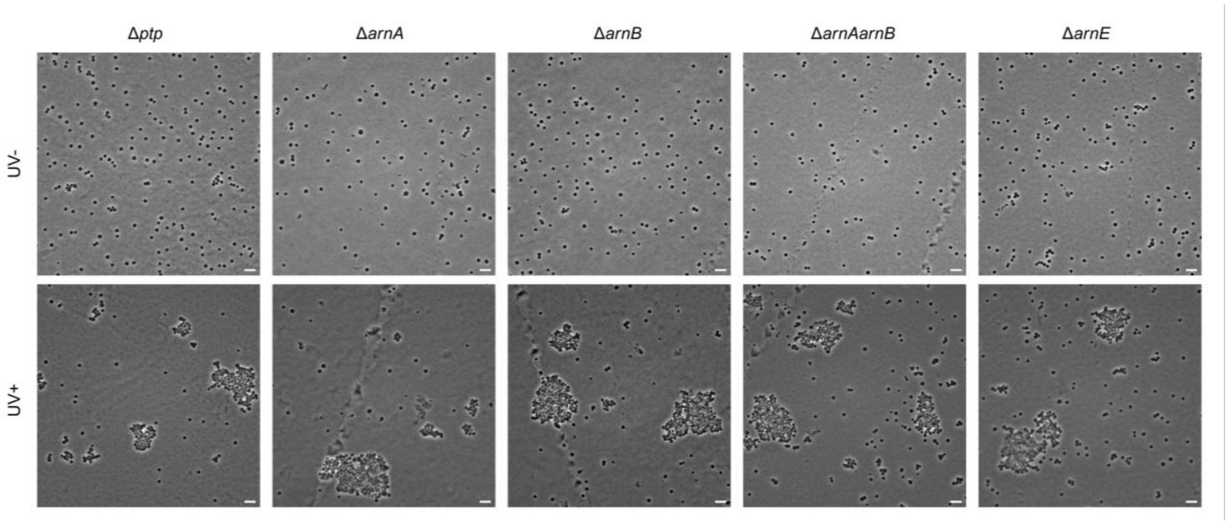


**Figure S4 UV-induced cell aggregation in *S. acidocaldarius* deletion strains.** Panels show representative microscopy images of *Δptp*, *ΔarnA*, *ΔarnB*, *ΔarnAΔarnB*, and *ΔarnE* cultures before (UV−) and after (UV+) UV irradiation, as indicated on the left of each row. Strain names are shown above the corresponding panels. Scale bar, 10 µm.
